# Mixtures of Intrinsically Disordered Neuronal Protein Tau and Anionic Liposomes Reveal Distinct Anionic Liposome-Tau Complexes Coexisting with Tau Liquid-Liquid Phase Separated Coacervates

**DOI:** 10.1101/2024.07.15.603342

**Authors:** Christine Tchounwou, Anjali J. Jobanputra, Dylan Lasher, Bretton J. Fletcher, Jorge Jacinto, Arjun Bhaduri, Rebecca L. Best, William S. Fisher, Kai K. Ewert, Youli Li, Stuart C. Feinstein, Cyrus R. Safinya

## Abstract

Tau, an intrinsically disordered neuronal protein and polyampholyte with an overall positive charge, is a microtubule (MT) associated protein, which binds to anionic domains of MTs and suppresses their dynamic instability. Aberrant tau-MT interactions are implicated in Alzheimer’s and other neurodegenerative diseases. Here, we studied the interactions between full length human protein tau and other negatively charged binding substrates, as revealed by differential-interference-contrast (DIC) and fluorescence microscopy. As a binding substrate, we chose anionic liposomes (ALs) containing either 1,2-dioleoyl-sn-glycero-3-phosphatidylserine (DOPS, −1e) or 1,2-dioleoyl-sn-glycero-3-phosphatidylglycerol (DOPG, −1e) mixed with zwitterionic 1,2-dioleoyl-sn-glycero-3-phosphatidylcholine (DOPC) to mimic anionic plasma membranes of axons where tau resides. At low salt concentrations (0 to 10 mM KCl or NaCl) with minimal charge screening, reaction mixtures of tau and ALs resulted in the formation of distinct states of AL-tau complexes coexisting with liquid-liquid phase separated tau self-coacervates arising from the polyampholytic nature of tau containing cationic and anionic domains. AL-tau complexes exhibited distinct types of morphologies. This included, large ≈20-30 micron tau-decorated giant vesicles with additional smaller liposomes with bound tau attached to the giant vesicles, and tau-mediated finite-size assemblies of small liposomes. As the ionic strength of the solution was increased to near and above physiological salt concentrations for 1:1 electrolytes (≈150 mM), AL-tau complexes remained stable while tau self-coacervate droplets were found to dissolve indicative of breaking of (anionic/cationic) electrostatic bonds between tau chains due to increased charge screening. The findings are consistent with the hypothesis that distinct cationic domains of tau may interact with anionic lipid domains of the lumen facing monolayer of the axon plasma membrane suggesting the possibility of transient yet robust interactions at physiologically relevant ionic strengths.

## INTRODUCTION

Tau, a microtubule-associate protein (MAP), is an intrinsically disordered overall positive polyampholyte containing positive and negative residues (Figure 1a). In mammals, there are six wild-type central nervous system (CNS) isoforms generated by alternative splicing of exons 2, 3 and 10.^1^ Tau’s N-terminal tail (NTT) is made up of a projection domain (PD) and proline rich region (PRR) (Fig. 1a). This is followed by the microtubule binding region (MTBR) and a carboxyl-terminal tail (CTT) (Figure 1b).^1–6^ The prevailing model is that tau binds the C-terminal tail of αβ-tubulin via the MTBR and possibly PRR regions enriched in positive residues.^1–6^ In our study we used the full-length isoform (4RL-tau, also known as 4R2N tau).

**Figure 1.**
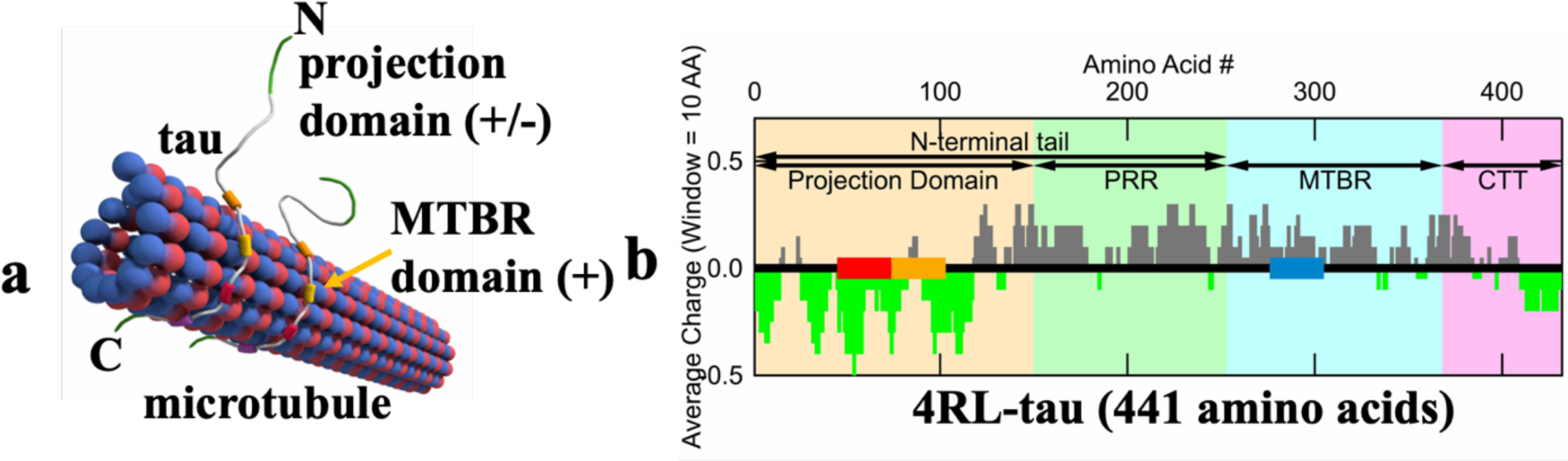
Illustration of tau binding to a microtubule and charge distribution of the full-length 4RL isoform of tau used in the study. (a) Tau binds to the surface of MT (red/blue) at the microtubule binding region (MTBR, teal background) and possibly the proline rich region (PRR, green background). The projection domain (PD, beige background) and C-terminal tail (background in pink) extend off the surface of the microtubule. (b) Charge distribution (averaged over 10 residues) of the primary sequence of full length 4RL-tau (negative charge in green, positive charge in grey). Data are from the National Center for Biotechnology Information Protein Database (accession number NP_005901.2).

In mature neurons, tau is confined primarily to the axonal compartment where it interacts with the microtubule (MT) cytoskeleton. MTs are made up of αβ-tubulin heterodimers with a net negative charge that self-organize into hollow nanotubes containing protofilaments (i.e. oligomers of αβ-tubulin) (Fig. 1b).^7–10^ MTs, in association with MAPs and other cellular proteins, play an important role in cell motility, establishing cell shape, intracellular trafficking, and chromosome segregation.^11,12^ Bare MTs are highly dynamic and stochastically switch between cycles of slow growth (polymerization) and rapid shrinkage (depolymerization) known as dynamic instability.^11,13–17^

While tau’s natural substrate is αβ-tubulin,^1–6,18–22^ it has additional roles in neurons many of which are not fully understood. In developing neurons, tau is associated with axon establishment and elongation, while in mature neurons, tau binds and stabilizes microtubules by suppressing dynamic instability (Figure 1a).^3–5,11,23–28^ This stabilization allows for proper trafficking of organelles along MTs in axons, a process known as axonal transport. However, tau dysfunction, such as thought to occur via hyperphosphorylation, leads to dissociation of tau from MT binding sites.^29^ The excess soluble tau forms phase separated fibers and tangles of fibers most likely driven by interactions with oppositely charged macromolecules such as RNA. These tau tangled states are known as neurofibrillary tangles and are very common in patients with Alzheimer’s Disease, Frontotemporal Dementia and Parkinsonism (FTDP-17) and several other dementia diseases.^30,31^

The experimental study described here was designed in order to identify interactions between full length 4RL-tau (Figure 1b) and other biologically relevant negatively charged binding substrates. A potential substrate for tau, in addition to tau’s natural substrate, namely, net negative αβ-tubulin whether in the MT lattice or not,^22,32^ may be the plasma membrane of axons where lipids with negative charge headgroups (e.g. phosphatidylserine or phosphatidylinositol-4,5 biphosphate) reside in the inner layer, facing the axonal cytosol.^33^ A previous study has reported interactions between tau’s overall negative projection domain and neural plasma membranes of PC12 cells composed of a complex mixture of lipids and proteins (i.e. raising the possibility of non-specific electrostatic interactions with cationic protein domains or specific tau/membrane-protein interactions).^34^ To model the negatively charged lipid domains of the plasma membrane, anionic liposomes were prepared and their reaction mixtures with 4RL-tau were investigated for identification of phase separated states resulting from the electrostatic interactions.

The initial studies involved tau-alone control experiments using 4RL-tau tagged with Alexa-546, visible in the RFP channel, in varying salt concentrations of KCl or NaCl, using both differential-interference-contrast (DIC) and fluorescence microscopy. Liquid-liquid phase separated (LLPS) tau self-coacervates were observed at low ionic strength. The LLPS arises from the polyampholytic nature of tau with cationic and anionic residues driving attractive electrostatic interactions between oppositely charged tau domains.^35^ The tau self-coacervates were observed to significantly weaken and effectively dissolve with increasing ionic strength. Previous observations at high ionic strength (i.e. in biological buffer) similarly found that tau coacervates can only retain their stability by the addition of a depletant resulting in osmotic crowding.^36–38^ Tau coacervates have been hypothesized to potentially be a precursor state to the neurofibrillary tangles in neurodegeneration.^35,39^

Similarly, the anionic liposome samples (with no added tau) consisting of zwitterionic 1,2-dioleoyl-sn-glycero-3-phosphatidylcholine (DOPC, charge = 0) mixed with either 1,2-dioleoyl-sn-glycero-3-phospho-L-serine (DOPS, charge =-1e) or 1,2-dioleoyl-sn-glycero-3-phospho-(1’-rac-glycerol) (DOPG, charge = −1e) and tagged with Cy5.5PE visible in the Cy5 Channel, exhibited morphologies (including giant multi-lamellar liposomes and small uni- and multi-lamellar vesicles), over the range of ionic strengths studied (0 mM to 300 mM, 1:1 electrolytes).

Optical imaging experiments of reaction mixtures of labeled tau and anionic liposomes (ALs), in varying salt concentrations, were conducted using DIC and fluorescence microscopy to elucidate the new phase separated states occurring in the reaction mixtures. Experiments with both the anionic liposomes unlabeled (50% DOPS/ 50%DOPC) or (50% DOPG/ 50%DOPC) and labeled (50% DOPS/ 49%DOPC/1% CY5.5 PE) or (50% DOPG/ 49%DOPC/1% CY5.5 PE) were performed with the labeled tau.

Distinct phase separated states were formed in the low salt concentration regime in mixtures with no added salt and with 10 mM KCl or NaCl. A large fraction of the reaction mixtures of tau and ALs consisted of AL-tau complexes. One distinct complex morphology included tau decorated giant uni- or multi-lamellar vesicles with additional smaller liposomes with bound tau, attached to the giant vesicle similar to colloidal nanoparticles bound to giant vesicles.^39^ A second common morphology was assemblies of small liposomes electrostatically bound together through the cationic domains of tau. These tau-mediated assemblies of small liposomes exhibited morphologies similar to vesosomes where for the latter system the vesicle assembly is achieved through encapsulation inside a larger unilamellar liposomsal compartment.^41,42^

A separate distinct state coexisting with AL-tau complexes at low ionic strength was the liquid-liquid phase separated tau coacervate, which appeared to contain no detectable lipid in fluorescence microscopy. Significantly, as a function of increasing NaCl or KCl concentrations from 0 to 300 mM, the AL-tau complexes remained stable, while the tau coacervate droplets were found to rapidly dissolve as the ionic strength approached physiological salt conditions (≈150mM 1:1 electrolyte). This behavior, also observed for tau-alone samples, is indicative of breaking of inter-tau (anionic/cationic) electrostatic bonds due to increased charge screening with increasing salt concentrations. The finding of stable AL-tau complexes at physiological (1:1 electrolyte) salt concentrations raises the possibility that the cationic domains of tau, either bound to MTs or upon unbinding from MTs, may form stable transient bonds with anionic domains of the inner layer of axonal plasma membrane.

It is noteworthy to point out that AL-tau complexes (or tau-lipoplexes), are analogous to well-known lipoplexes,^43^ consisting of complexes of cationic liposomes (CLs) and oppositely charged nucleic acids (DNA, gene silencing siRNA), used in delivery applications.^44–46^ More broadly, CL– DNA, CL-siRNA and CL-anionic peptide lipoplexes exhibit distinct structures as revealed by synchrotron small-angle X-ray scattering. These include, the lamellar L_α_^C^ phase and PEGylated lamellar lipoplex nanoparticles with alternating lipid bilayers and DNA monolayers,^47–50^ and inverse hexagonal H_II_^C^ and hexagonal H_I_^C^ phases with DNA inserted in inverse cylindrical micelles or surrounded by cylindrical micelles, respectively.^51–54^ For short double stranded siRNA, CL–siRNA complexes may further form inverse cubic phases with siRNA confined to nanoscale water channels.^55–58^ The structures of recently developed mRNA-lipid nanoparticles (LNPs) used in vaccines, similarly reveal a disordered inverse hexagonal H_II_^c^ and cubic phases.^59,60^

## MATERIALS AND METHODS

### Tau Purification

The expression vector (pRK172) encoding full-length (4RL) tau with ampicillin resistance was gifted by Dr. Kenneth Kosik (University of California, Santa Barbara). Tau plasmids were transformed into BL21(DE3) competent cells and streaked on Luria broth agar (5g tryptone, 2.5g yeast extract, and 5g NaCl in DI water) plates containing 0.1% ampicillin before a ∼14hr incubation at 37°C. A single colony was inoculated in 250 mL of Luria broth overnight (18hr) at 37°C followed by a 24h incubation in 6L of auto-induction media (10g tryptone, 5g yeast extract, 0.5g dextrose, 2g α-D-lactose, and 5 mL of glycerol per liter of DI water with 25mM NaHPO_4_, 25mM KH_2_PO_4_, 50mM NH_4_Cl, 5mM Na_2_SO_4_ dissolved). Bacteria were harvested for lysis by centrifugation (5,000rpm for 15min, GS-3 Rotor) in a Sorvall RC-5B Plus centrifuge held between 4 to 10 °C. Pelleted bacteria were resuspended in BRB80 buffer (80mM PIPES, 1mM EGTA and 1mM MgSO_4_) pH 6.8 which includes ∼120mM NaOH (to obtain the correct pH) as well as 0.1% βME and 0.5% AEBSF (a protease inhibitor). Following, cells were lysed in a pressurized French press cell three times at 1200 PSI, boiled and cooled for 10 min each, and centrifuged at 13,000rpm in a SS-34 Rotor for 40 min to pellet the cell debris. The supernatant was collected and passed through a phosphocellulose (P11, Whatman) ion-exchange column to initiate isolation of tau by increasing concentrations of (NH_4_)_2_SO_4_ (50mM, 150mM, and 250mM) in BRB80 for elution. The tau fractions, confirmed by SDS-PAGE, were pooled and the buffer concentration of (NH_4_)_2_SO_4_ increased to 1.25M. The sample was then passed through a hydrophobic interaction chromatography column (HisTrap Phenyl HP, GE Healthcare) and washed with decreasing concentrations (1.1 M – 50 mM) of (NH_4_)_2_SO_4_ in BRB80 to elute tau. Tau containing fractions were identified again through SDS-PAGE, pooled, concentrated, and buffer exchanged to fresh BRB80 without (NH_4_)_2_SO_4_ using an Amicon Ultra-15 Centrifugal Filter with MWCO = 10,000 (EMD Millipore, Darmstadt, Germany) and successive 40min centrifugation cycles at 4,000 rpm. Purified tau stocks were stored at −80 °C until needed for experiments. The concentration of tau was determined by SDS-PAGE slope comparison with a tau mass spectrometry standard.

### Tau Labeling

To attach a fluorophore to purified tau stocks, tau’s concentration was confirmed between 50-100µM in BRB80 (pH ∼7.0) and thawed in 100-500µL batches. Next a 10-molar excess of βME was added and the solution was left to sit for 30 minutes at room temperature to reduce tau’s disulfide bonds. βME was dialyzed away using a Zeba Spin desalting column (MWCO 7,000, Thermo Fisher Scientific). A 1-10mM stock solution of the dye was then made followed by adding the dye to the tau solution dropwise. In this case, Alexa Fluor-488 C5 maleimide or Alexa Fluor-546 C5 maleimide (Thermo Fisher Scientific) was added depending on experimental parameters. After two hours at room temperature, a 10-molar excess of βME was added and a gel filtration column (Sephadex G-25 Column, Sigma Aldrich) was performed in BRB80 to separate the conjugate. Labeled tau was then stored at – 80 °C until experimental use.

### Liposome Preparation

In a small glass vial, stock solutions of each lipid component stored in chloroform purchased from Avanti Polar Lipids, Inc were mixed in chloroform:methanol (3:1, v/v) at a total molar concentration dependent on the molar charge ratio (ρ) of lipid/tau =1. For the experiments outlined, liposomes were formulated with lipids of negative charge,1,2-dioleoyl-sn-glycero-3-phosphatidylserine (DOPS, −1e) or 1,2-dioleoyl-sn-glycero-3-phosphatidylglycerol (DOPG, −1e), mixed with zwitterionic lipids, 1,2-dioleoyl-sn-glycero-3-phosphatidylcholine (DOPC) and some with the fluorophore 1,2-dioleoyl-sn-glycero-3-phosphoethanolamine-N- (Cyanine 5.5) as well. The ratios were based on the final desired concentration and molar composition of each lipid. Liposomes of various molar ratios were formulated including (50% DOPS/ 50% DOPC), (50% DOPG/ 50% DOPC), (50% DOPS/49% DOPC/1% CY5.5 PE), and (50% DOPG/ 49% DOPC/1% CY5.5 PE). Lipid mixtures in chloroform:methanol were exposed to a steady stream of nitrogen for 10 min to evaporate the organic solvent and then further dried overnight in a vacuum (rotary vane pump). Dehydrated lipid films were then rehydrated with high-resistivity water (18.2 MΩ cm) determined by the final desired concentration and incubated overnight at 37 °C to form unsonicated liposomes. After incubation, the liposomes were stored at 4 °C until needed for experiments.

### Differential-Interference-Contrast (DIC) Microscopy

For microscopy experiments, tau tagged with Alexa Fluor-546 (excitation/emission: 561nm/572nm) was thawed over ice and solutions of 0.73 µg/µL tau (1.59×10^−14^ M) in varying salt solutions (0-300mM) were created in separate Eppendorf tubes at room temperature. 3µL of each solution was placed on microscope slides, along with a coverslip and parafilm to maintain a seal. The samples were immediately imaged on the Nikon Eclipse Ti2 Microscope at 20x, 40x, or 60x objective with the addition of a 1.5x lens. Liposomes were retrieved from 4°C and mixed into varying salt concentrations (10-300mM) in separate Eppendorf tubes at room temperature. Molar compositions of liposomes varied based on the experimental objective. Microscope slides were created and samples were imaged in the same format as aforementioned. Liposome samples prepared at 60µM concentration were mixed with 0.73 µg/µL tau (1.59×10^−14^ M) at a volume charge density (ρ) of 1 in an Eppendorf tube in varying salt solutions (0-300mM). Similarly, two different types of tau tagged with fluorophores were used depending on the liposome mixture. If liposomes were tagged with a fluorophore (Cy5.5 PE lipid - excitation/emission 683nm/703nm), then tau tagged with Alexa-488 (Alexa 488 excitation/emission: 488nm/496nm), was used whereas Alexa-546 (Alexa 546 excitation/emission: 561nm/572nm) tau was used when lipids were not labeled. The sample solutions of the liposome-tau mixture were stored at room temperature for the duration of the experiment. The microscope slide containing the liposome-tau mixture was created and imaged using the previous protocol for imaging tau alone.

### Fluorescence Microscopy

Microscope slides were prepared in the same method outlined for DIC and imaged with the Nikon Eclipse Ti2 in fluorescence mode. To observe the liposomes present in the sample, the Cy5 (Cyanine 5) filter was used (Cy5.5 PE lipid - excitation/emission 683nm/703nm). To observe tau present, either the GFP (Green fluorescent protein) channel (Alexa Fluor 488; excitation/emission: 488nm/496nm) or the RFP (Green fluorescent protein) channel was used (Alexa Fluor 546; excitation/emission: 561nm/572nm). Again, the samples were viewed using the 20x, 40x, or 60x objective with the addition of a 1.5x lens.

## RESULTS AND DISCUSSION

### Salt-Dependent Behavior of 4RL Tau Self-Coacervates

In an aqueous system at very low ionic strength, 4RL-tau composed of 441 amino acids was found to form stable liquid-liquid phase separated (LLPS) droplets as depicted in the cartoon in figure 2a. Figure 2 (b-f) shows differential-interference-contrast (DIC) micrographs of tau self-coacervates as a function of increasing ionic strength between 0 mM and 300 mM KCl. Here, we used full length 4RL-tau tagged with Alexa-546 visible in the RFP channel, in order to carry out near simultaneous DIC and fluorescence experiments on the same samples. At 0mM (deionized Milli-Q; pH 6.998) and 10 mM KCl, there is an abundance of tau droplets as shown in figure 2 (b, c and zoomed-in section). The observation of stable droplets is due to protein tau’s inherent polyampholyte nature leading to intermolecular electrostatic interactions between tau chains and self-coacervation (figure 2a).^35^ To form tau self-coacervates, the cationic sections of 4RL-tau (figure 1a), made up of the PRR (amino acid 152-240) and the MTBR (amino acid 241-368) domains containing exon 10 and thus all 4 repeats, must have favorable electrostatic interactions with the anionic domains at the N- and C-terminals made up of the PD (amino acid 1-151) containing the inclusion of exon 2 and 3 and the CTT (amino acid 369-441).

**Figure 2.**
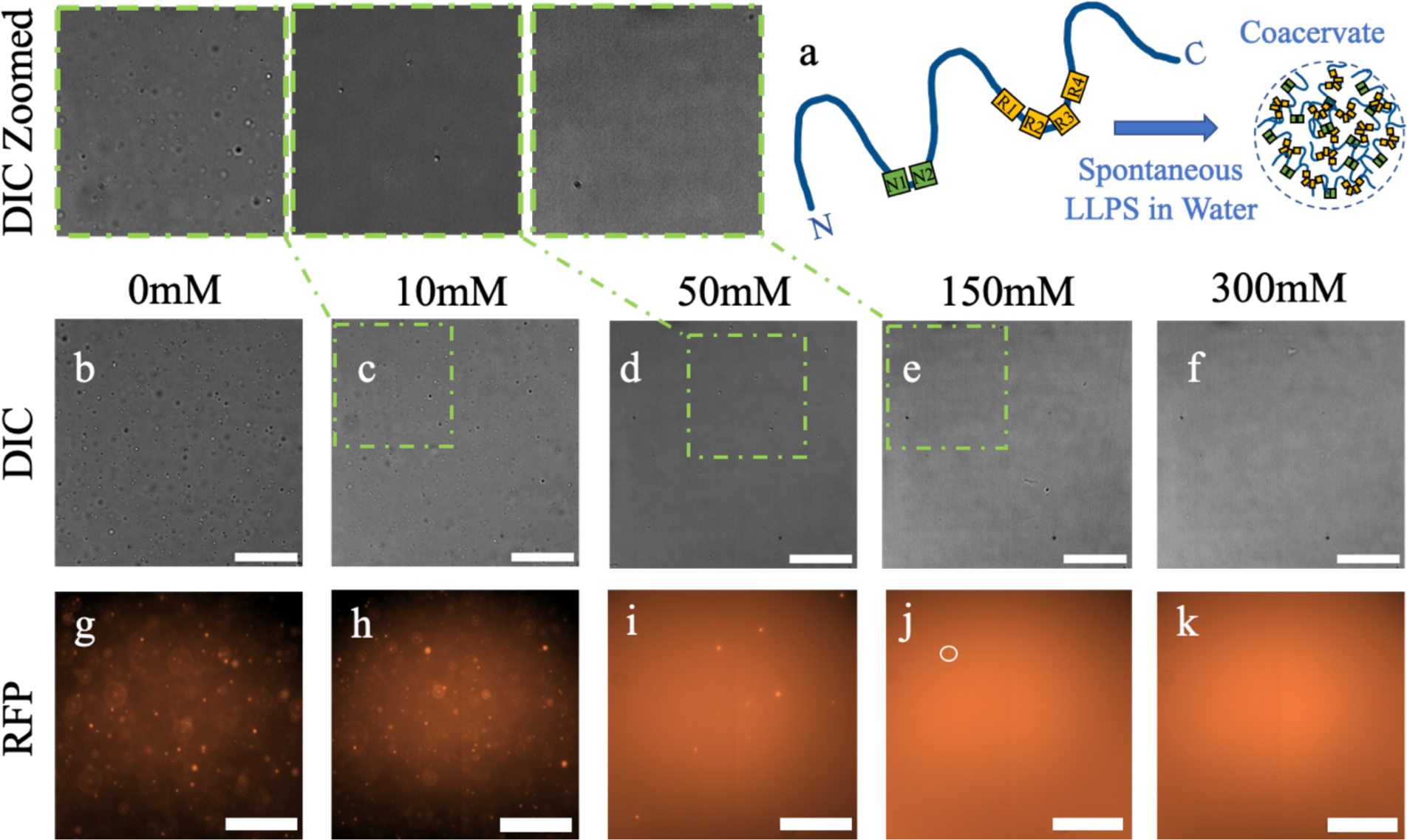
Attractive electrostatic interactions stabilizing tau self-coacervates break down near physiological salt concentration. (a) Cartoon of 4RL-tau polyampholyte undergoing intermolecular electrostatic interactions between anionic and cationic domains of tau chains to create a tau coacervate in deionized water. (b-f) Differential interference contrast (DIC) microscopy images of full length 4RL-tau tagged with Alexa-546 in increasing KCl concentrations (0-300mM). Zoomed-in inset of corresponding DIC images (c-e) linked by dashed line scaled by a factor of 2. (g-k) Fluorescent microscopy of full length 4RL-tau tagged with Alexa-546 visible in the RFP channel (Alexa 546 excitation/emission: 561nm/572nm) in increasing KCl concentrations (0-300mM). All scale bars = 50µm.

A significant contrasting behavior is observed with increasing salt concentration. At 50mM KCl (Figure 2d and zoomed-in section) there are far fewer particles, and little to none at 150mM near physiological concentrations (Figure 2e and zoomed-in section) and beyond at 300 mM (Fig. 2f). The breakdown of tau self-coacervates is more clearly seen in fluorescence images of the same particles with 4RL-tau tagged with Alexa-546 (Figure 2g-k). At 0mM and 10mM KCl (Figure 2g,h), there is again an abundance of particles and little diffuse background indicative of free tau. In contrast, at 50mM (figure 2i), there are far fewer particles and the background fluorescence is increasing, indicating a coexistence of tau coacervates and tau dispersed in solution. At 150mM KCl (figure 2j), near physiological salt concentrations, there are almost no particles and the background is mostly diffused, showing that the majority of tau is no longer in the LLPS droplet state. An occasional rare coacervate in the high salt regime is indicated by the white circle in figure 2j.

The observation of dissolution of tau self-coacervate droplets, in going from 10 mM to 150 mM KCl, implies a significant decrease in the attractive electrostatic forces between anionic and cationic domains of tau likely due to enhanced charge screening and a decrease in the Debye screening length from λ_D_ ≈3.04 nm to ≈0.785 nm. This observation is consistent with previous studies at high salt where it was found that the tau self-coacervates can retain their stability only with the addition of a depletant such as poly-(ethylene-oxide) (PEO), giving rise to PEO-induced depletion attraction between tau chains, phase separation and droplet formation.^36–38^

### Anionic Liposomes at Low and High Salt Concentrations

Figure 3a depicts the chemical structures of the lipid components used when forming the anionic liposomes: DOPS (charge = −1e), DOPG (charge = −1e), DOPC (C18:1, zwitterion, charge = 0) and a small amount of Cy5.5-PE (C18:1) to add a fluorescent tag. Due to lipids poor solubility in water, they are mixed in an organic solvent, allowing each individual lipid component to easily dissolve (see Methods). This creates liposomes made up of a uniform mixture. The anionic liposomes had a molar composition of either (50% DOPG/ 49% DOPC/1% CY5.5 PE) or (50% DOPS/ 49% DOPC/1% CY5.5 PE).

**Figure 3.**
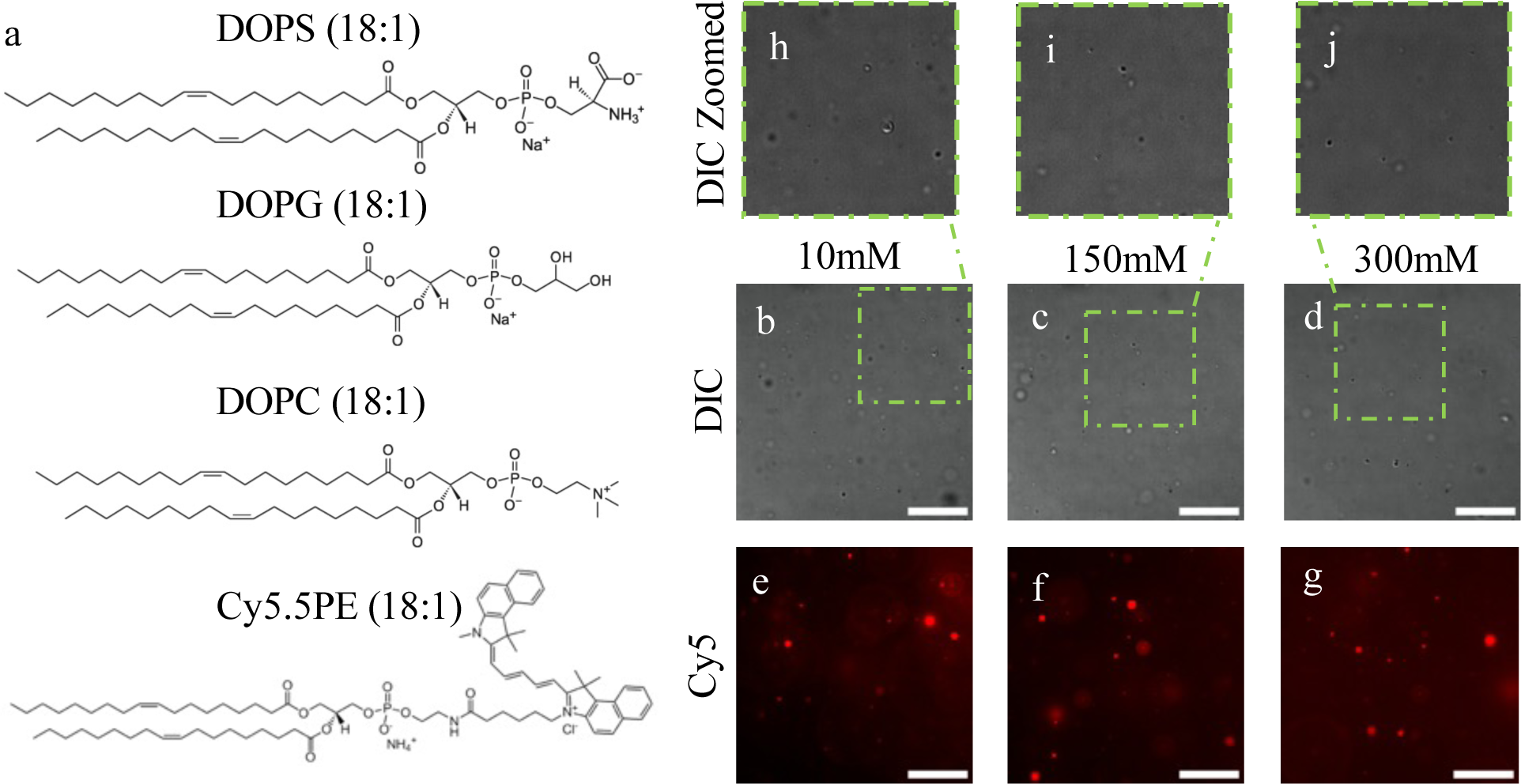
Observation of anionic liposomes at low and high salt conditions. (a) Chemical structures of DOPS (C18:1, charge = −1e), DOPG (C18:1, charge = −1e), DOPC (C18:1, zwitterion) and Cy5.5-PE (C18:1) lipids used to make anionic liposomes for experiments.^61^ (b-d) DIC microscopy images of unsonicated anionic liposomes tagged with Cy5.5-PE (50 mol% DOPS/ 49 mol% DOPC/1 mol% CY5.5 PE) in increasing KCl concentrations (10-300 mM). (e-g) Fluorescent microscopy of unsonicated anionic liposomes tagged with Cy5.5-PE (50 mol% DOPS/ 49 mol% DOPC/1 mol% CY5.5 PE) visible in the Cy5 channel (Cy5.5 PE lipid - excitation/emission 683nm/703nm) in increasing KCl concentrations (10-300 mM). (h-j) Zoomed-in inset of corresponding DIC images linked by dashed line scaled by a factor of 2. All scale bars = 50µm.

Figure 3 (b-d) shows DIC images of anionic liposomes as a function of increasing ionic strength between 10 mM and 300 mM KCl. The number of liposoms remain nearly constant between 10mM (figure 3b) and 300mM (figure 3d), suggesting that the anionic liposomes are stable and retain their structure under both low and high ionic strengths. This is further confirmed in fluorescence images of the same particles seen in DIC (figure 2e-f). Both the background and the concentration of particles appear similar between 10mM KCl (figure 3e) and 300mM KCl (figure 3g).

### Phase Separated States in Mixtures of Anionic Liposomes and Tau in Deionized Water

To elucidate the types of distinct states/structures formed when anionic liposomes (ALs) interact with polyampholyte tau in the regime where electrostatic interactions are the strongest, we studied a reaction mixture of tau and liposomes in deionized water containing either DOPS (−1e) or DOPG (−1e). Similar behavior was found for studies at 10 mM KCl. Phosphotidylserine (PS) based anionic lipids are among the primary lipids imparting negative charge to mammalian plasma membranes, whereas, lipids with phosphotidylglycerol (PG) headgroups constitute a second class of anionic lipids occurring at a lower content (1-2%) compared to PS-based anionic lipids.^12^ Figure 4a shows optical fluorescent images of ALs (50% DOPG/ 49%DOPC and 1% CY5.5-PE fluorescent in the CY5 (red) channel) mixed in deionized water with 4RL-tau tagged with Alexa- 488, fluorescent in the GFP (green) channel at a lipid/tau molar charge ratio ρ=1. A z-stack of images in the x-y plane (with merged GFP and CY5 channels), allows for visualization of the 3D structures of the particles in the x-y, y-z, and x-z planes and reveals three distinct phase separated states.

**Figure 4.**
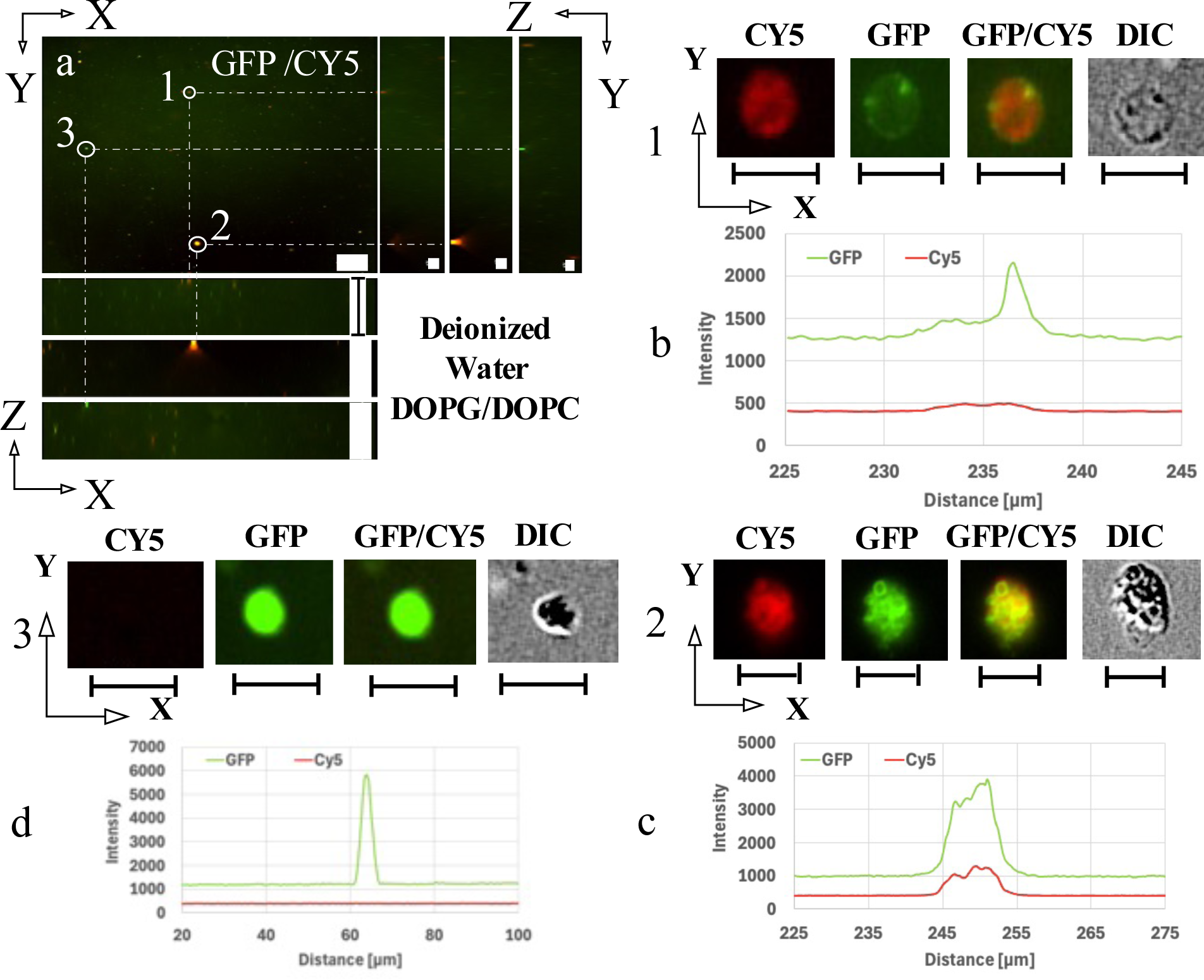
Fluorescent microscopy images of anionic liposomes made of 50 mol% DOPG/ 49 mol%DOPC/1 mol% CY5.5 PE interacting with 4RL-tau in deionized water. The lipid channel (Cy5) is shown in RED (excitation/emission 683nm/703nm) and 4RL-tau tagged with Alexa-488, is shown in the GFP channel (green) (excitation/emission 488nm/496nm). (a) A 3D Z-stack image series showing both GFP and Cy5 channels. The main image frame (top left panel) shows the X-Y section view at a chosen Z. The projection to the right shows the Y-Z side views slicing along the vertical dotted lines of three circled objects 1-3. The bottom panel shows the X-Z slices along the horizontal dotted lines. The X-Z and Y-Z plane side views of each circled object are indicated by the end of the dotted lines in each direction. (Scale bars correspond to 50µm in the X-Y plane, 200µm in X-Z and 20µm in Y-Z planes, respectively) (b) Detail view of particle 1: anionic liposome-4RL tau complex consisting of a tau coated giant multilamellar vesicle. Top panel from left to right: magnified fluorescent (Cy5, GFP and GFP/Cy5 overlay) and DIC images of particle 1. Smaller liposomes with bound tau are also seen attached to the giant vesicle. Bottom panel: fluorescent intensity profiles of particle 1 in Cy5 and GFP channels as a function of X. (c) Detail view of particle 2: an anionic liposome-4RL tau complex, consisting of tau-mediated assembly of small liposomes, shown in magnified fluorescent (Cy5, GFP and GFP/Cy5 overlay) and DIC images, and fluorescent intensity traces as a function of X. (d) Detail view of particle 3: a tau self-coacervate (lacking lipid) shown in magnified fluorescent (Cy5, GFP and GFP/Cy5 overlay) and DIC images, and fluorescent intensity traces as a function of X. Scales bar in (b-d) = 10µm.

Particles labeled 1 and 2 in figure 4a, visible in both GFP and CY5 channels, are characterized as anionic liposome-tau complexes. In the first AL-tau complex state (labeled “1” in figure 4a with expanded views in the top four panels in figure 4b), 4RL-tau (seen in the GFP channel) coats the perimeter of a giant multilamellar vesicle. The complex further includes smaller liposomes with bound tau attached to the giant vesicle (i.e. smaller vesicles seen in the Cy5 (lipid) channel and the corresponding tau clumps in the GFP channels). The green (tau) and red (lipid) intensity scans through the AL-tau complex (Figure 4b, lower panel), are consistent with the described structure where you can see the overall “U” shapes and the asymmetric distribution of tau (green line) around the giant multilamellar vesicle (red line).

A second distinct AL-tau complex state observed (labeled “2” in figure 4a with expanded views in the top four panels in figure 4c), consists of a tau-mediated assembly of small liposomes facilitated through the cationic domains of 4RL-tau sandwiched between the surface of the vesicles. This is confirmed in the particle images where individual spheres bundled together are observed in the GFP, CY5, and GFP/CY5 merged channels. The green (tau) and red (lipid) intensity profiles in figure 4c show multiple peaks for both GFP and Cy5 channels consistent with an assembly of tau-coated vesicles. The morphology of the tau-mediated assembly of small liposomes is similar to that of vesosomes used in drug delivery applications. However, the small neutral vesicles in vesosomes are brought together from confinement forces within an outer membrane bilayer.^41,42^

A third distinct phase separated state further appears in mixtures of ALs and tau. This particle labeled “3” in figure 4a (with expanded views in the top four panels in figure 4d) is an example of self-coacervation of polyampholyte tau through intermolecular electrostatic interactions between the anionic and cationic domains (figures 1 and 2a). As expected for a liquid-liquid phase separated state, the particle, which is seen in the GFP channel but not the CY5, is spherical. Consistently, the green (tau) and red (lipid) intensity profiles in figure 4d display a highly symmetrical tau peak indicative of a liquid coacervate with no evidence of any lipid component.

Figure 5 displays fluorescent images similar to those of figure 4, but instead examines the interaction of 4RL-tau (tagged with Alexa-488) and anionic liposomes with composition (50% DOPS/ 49%DOPC/1% CY5.5 PE) in deionized water, where DOPG is replaced by DOPS at a lipid/tau molar charge ratio ρ=1. Three particles with different structures are identified in Figure 5a displaying images in the x-y, x-z, and y-z planes with the GFP and CY5 channels merged. The expanded images of particle 1 (top panels (CY5, GFP, GFP/CY5, DIC) in figure 5b) display another example of 4RL-tau decorating the perimeter of a giant multilamellar vesicle with the same “U” shape green and red intensity profiles (figure 5b). For this particle, fewer small liposome-tau particles are observed to be attached to the giant vesicle, compared to particle 1 from figure 4. Particle 2 (top panels (CY5, GFP, GFP/CY5, DIC) in figure 5c) is another example of an assembly of small liposomes fused together by tau.

**Figure 5.**
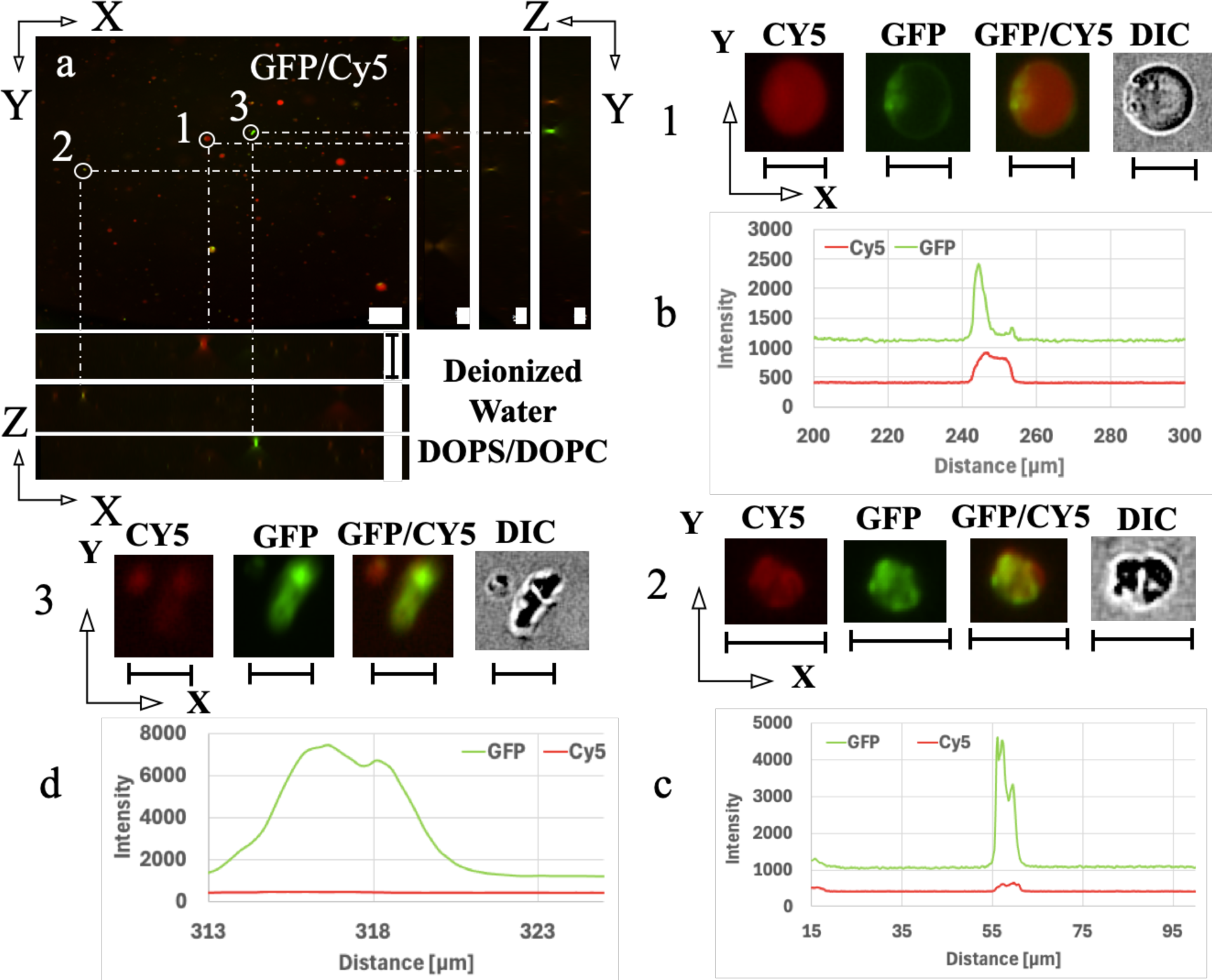
Fluorescent microscopy images of mixtures of anionic liposomes made of 50 mol% DOPS/49 mol% DOPC/1 mol% CY5.5 PE and 4RL-tau in deionized water. The lipid channel is shown in the RED Cy5 channel (excitation/emission 683nm/703nm) and 4RL tau tagged with Alexa-488 is shown in green in the GFP channel (excitation/emission 488nm/496nm). (a) A Z-stack image series showing both GFP and Cy5 channels. The top left frame shows the X-Y plane view at a chosen Z. The three panels to the right show the Y-Z slices along vertical dotted lines marking three circled objects 1-3. The bottom panels are the X-Z side views (slices) along the horizontal dotted lines of circles objects 1-3. The Z-X and Y-Z plane side views of each circled object are indicated by the end of the dotted lines in each direction. (Scale bars correspond to 50µm in the X-Y, 200µm in X-Z and 20µm in Y-Z planes, respectively) (b) Detail view of particle 1: a giant anionic liposome-4RL-tau complex with a few small liposome-tau particles attached to the giant vesicle. Top panel from left to right: magnified fluorescent (Cy5, GFP and GFP/Cy5 overlay) and DIC images of particle 1. Bottom panel: fluorescent intensity profiles of particle 1 in Cy5/GFP channels as a function of X. (c) Detail view of particle 2: an anionic liposome-4RL tau complex (assembly of small liposomes glued together by tau) shown in magnified fluorescent (Cy5, GFP and GFP/Cy5 overlay) and DIC images, and fluorescent intensity traces as a function of X. (d) Detail view of particle 3 shown in magnified fluorescent (Cy5, GFP and GFP/Cy5 overlay) and DIC images, and fluorescent intensity traces GFP (tau) and CY5 (lipid) as a function of X. As discussed in the text, the object consists of a tau coacervate (dense semi-spherical particle in the top right in the GFP-tau channel) bound to an anionic liposome-tau complex consisting of a non-spherical unilamellar vesicle (dimly visible in Cy5 channel) coated by tau with hollow regions typical of liposomes visible in the GFP channel. Scales bar in (b-d) = 10µm.

In contrast, particle 3 (top panels (CY5, GFP, GFP/CY5, DIC) in figure 5d) displays a less commonly observed structure in mixtures of ALs and tau. Here, a tau coacervate (dense semi-spherical particle in the top right in the GFP-tau channel) appears to be bound to a AL-tau complex consisting of a non-spherical liposome coated by tau with hollow regions typical of liposomes visible in the GFP channel. The GFP (tau) intensity profile (lower panel, figure 5d), which displays multiple peaks is supportive of a tau-coacervate interacting with an AL-tau complex. In contrast to the strong GFP intensity profile, there is a lightly visible non-spherical vesicle shape in the Cy5 channel that is not picked up in the red intensity profile (figure 5d). Intensity profiles taken in the x-y plane through different sections of the non-spherical shaped particle (with a concave distortion) in the Cy5 channel are similarly flat (Supporting Information, figure S1). This is consistent with the non-spherical vesicle being uni-lamellar with relatively few fluorescent lipids sampled in the line scans.

### Comparison of Phase Separated States in Mixtures of Anionic Liposomes and Tau in Low and High Salt Concentrations

Figure 6 compares mixtures of unlabeled anionic liposomes (50 mol% DOPG/ 50 mol% DOPC) and 4RL-tau tagged with Alexa-546, fluorescent in the RFP channel, at low and high salt concentrations in DIC and the corresponding fluorescent images. Two different salts, KCl and NaCl were used near physiological 1:1 electrolyte concentrations. Figures 6a, 6c, 6e, and 6g show the interactions at a low ionic strength of 5mM, while figures 6b, 6d, 6f, and 6h were performed at a high ionic strength of 150mM and the particles found in the RFP channel were circled. There is a clear decrease in the number of particles in the DIC micrographs as the mixtures transitioned from low (5 mM, figures 6a, 6e) to high salt (150 mM, figures 6b, 6f). Remarkably, the corresponding fluorescence images show that the decrease in particle density is accompanied by an increase in background fluorescence due to free labeled tau as the mixtures transition from low (figures 6c, 6g) to high salt (figures 6d, 6h).

**Figure 6.**
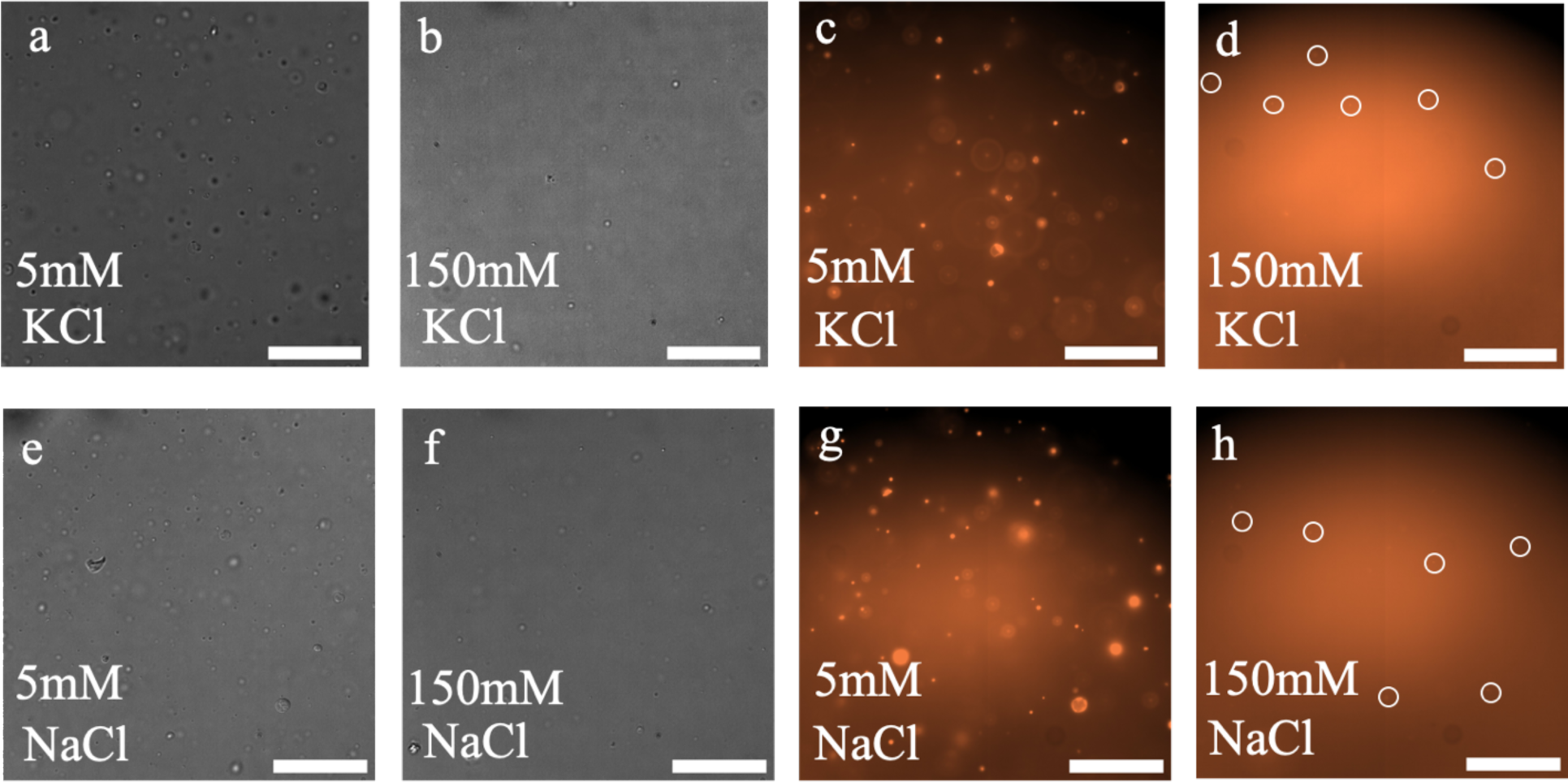
Attractive electrostatic interactions in tau coacervates are weaker than in anionic liposome-tau complexes. (a, b) DIC microscopy images of full length 4RL-tau tagged with Alexa-546 and unlabeled anionic lipids (50 mol% DOPG/ 50 mol% DOPC) in low (5 mM) and high (150 mM) KCl concentrations. (c, d) fluorescent microscopy images in the RFP channel corresponding to (a,b) showing full length 4RL-tau tagged with Alexa-546 (excitation/emission 561nm/572nm). [50µm scale bar] (e, f) DIC microscopy images of full length 4RL-tau tagged with Alexa-546 and unlabeled anionic lipids (50 mol% DOPG/ 50 mol% DOPC) in low (5 mM) and high (150 mM) NaCl concentrations. (g, h) RFP fluorescent images corresponding to DIC images shown in (e, f). [50µm scale bar]

At the low ionic strengths, the particles observed in DIC and fluorescence micrographs from labeled tau (figures 6a, 6c, 6e, and 6g) include both tau coacervates and AL-tau complexes. For tau alone preparations at increasing salt concentrations (figure 2), we observed that tau self-coacervates dissolved at concentrations between 100 mM and 150 mM (with very few coacervates remaining at 150 mM) leading to the observed increase in fluorescence background due to free tau. Thus, the observed particles at 150 mM NaCl or KCl (figures 6b, 6d, 6f, and 6h), mainly correspond to AL-tau complexes, which remain stable while coacervate droplets disperse into free tau.

Another less common but noteworthy structure, observed in mixtures of ALs (50% DOPS/ 49%DOPC/1% CY5.5 PE) and tau in 100 mM KCl, consisted of AL-tau complexes displaying an assembly of tau-free “bare” liposomes trapped inside a giant multilamellar vesicle (i.e. similar to vesosomes^40,41^) with tau decorating the outside perimeter. This is shown in figure 7 (labeled “1” in the merged GFP and CY5 channels in figure 7a with expanded views in the top four (CY5, GFP, GFP/CY5, DIC) panels in figure 7b). The lack of tau associated with the internal vesicles can be seen in comparing the GFP (tau) and CY5 (lipid) channels, where tau coats the giant vesicle surface and also a few much smaller liposomes bound to the surface of the giant vesicle. The GFP (tau) and red (lipid) intensity profiles (lower panel, figure 7b) are also consistent with the observed structures in the GFP, CY5, and GFP/CY5 channels. No tau self-coacervates (which are rare at these salt concentrations) were found in this mixture at 100 mM and the background GFP fluorescence indicative of dispersed tau is significantly higher compared to mixtures at 5 mM KCl. It is important to note that at this high salt concentration no tau-free liposomes were observed; instead, they only existed as AL-tau complexes.

**Figure 7.**
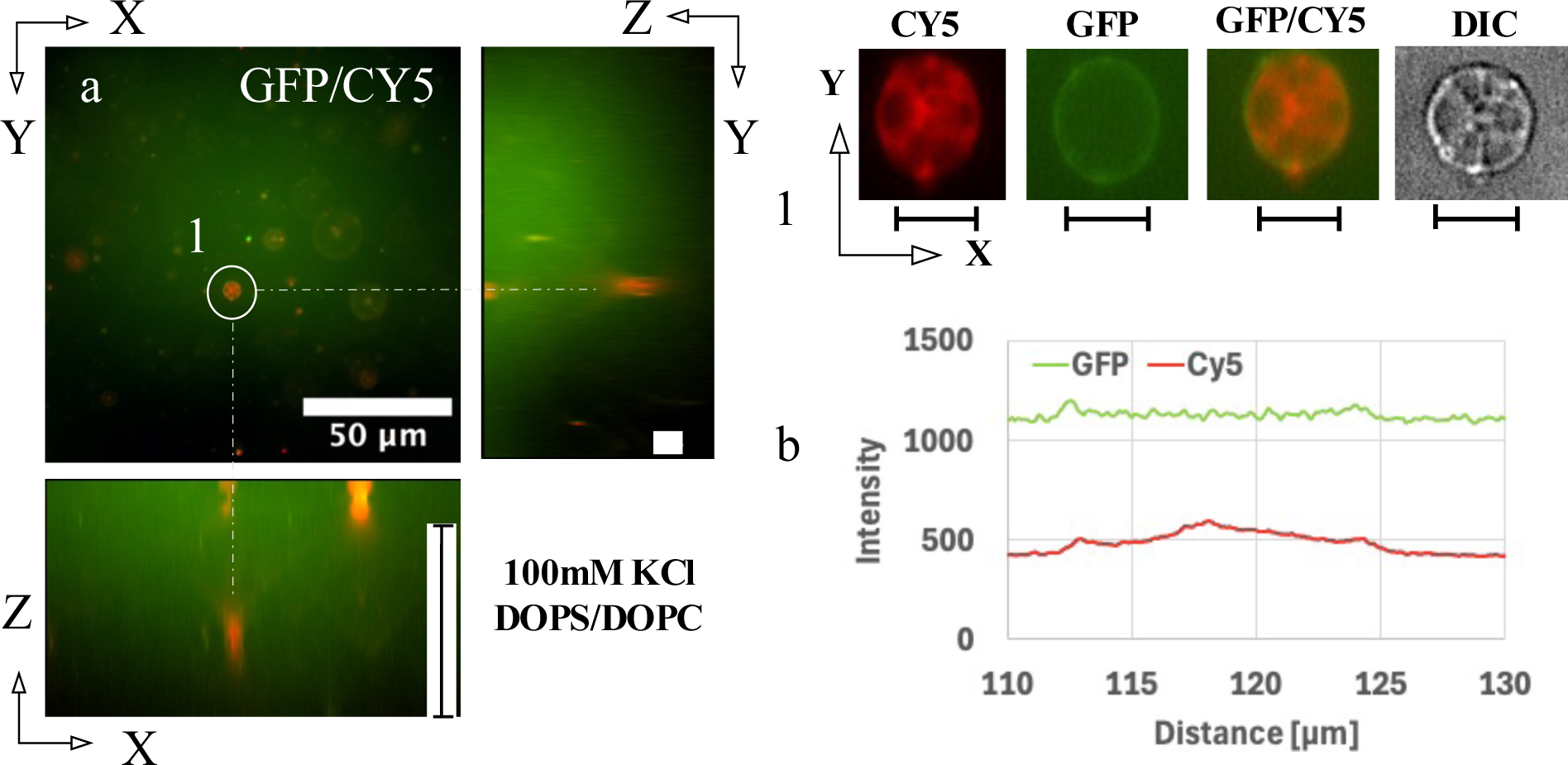
Fluorescent microscopy images of mixtures of anionic liposomes and 4RL tau in 100mM KCl. The anionic liposomes made from 50 mol% DOPS/ 49 mol% DOPC/1 mol% CY5.5 PE are visible in the Cy5 channel (red) (Cy5.5 PE lipid - excitation/emission 683nm/703nm) and 4RL-tau tagged with Alexa-488 (Alexa 488 excitation/emission 488nm/496nm) are visible in the GFP channel (green). (a) A Z- stack image showing both GFP and Cy5 channels. The top left frame shows the X-Y section view at a chosen Z. The panels to the right show the Y-Z slice along the vertical dotted line marking the circled object 1. The bottom panel shows the X-Z section along the horizontal dotted line of object 1. (Scale bars correspond to 50µm in the X-Y, 200µm in X-Z and 20µm in Y-Z planes, respectively) (b) Detail view of particle 1: a giant anionic liposome-4RL tau complex consisting of an assembly of bare liposomes trapped inside a giant multilamellar vesicle. As discussed in the text, tau is observed to coat the giant vesicle surface and also a few small liposomes bound to the giant vesicle. Top panel from left to right: magnified fluorescent (Cy5, GFP and GFP/Cy5 overlay) and DIC images of particle 1. Bottom panel: fluorescent intensity profiles of particle 1 in Cy5/GFP channels as a function of X. [10µm scale bars]

Taken together, our observation indicates that the electrostatic interactions between the cationic domains of tau and anionic membranes are significantly stronger than the electrostatic interactions stabilizing tau self-coacervate droplets, which are found to dissolve and disperse tau with increasing (1:1 electrolyte) salt concentrations.

## CONCLUSION

Intrinsically disordered protein tau, an overall positive polyampholyte, is a microtubule (MT)- associated protein localized to axons in vertebrate neurons. Tau regulates MT dynamic instability and aberrant tau-MT interactions is implicated in Alzheimer’s and other neurodegenerative diseases. While there is incomplete understanding of the mechanisms driving neurodegenerative pathology, some evidence exists that aside from tubulin and microtubules (tau’s best studied substrates) tau’s N-terminal projection domain can interact with other substrates including neural plasma membranes in PC12 cells, which contain a heterogeneous mixture of lipids and proteins (i.e. where the protein components may have specific or non-specific charge interactions with tau’s overall negative projection domain).^34^ However, the extent and conditions of these interactions are not understood.

We investigated the structural implications of the electrostatic associations of overall positive human 4RL-tau with “protein-free” anionic liposomes, containing either DOPS (charge = −1e) or DOPG (charge = −1e) mixed with zwitterionic DOPC, to mimic anionic lipid domains of plasma membranes of axons where tau resides. Using differential-interference-contrast and fluorescence microscopy, we studied the phase behavior of mixtures of 4RL-tau and anionic liposomes (ALs) at low and high salt concentrations at lipid/tau molar charge ratio equal to one. At low ionic strengths with minimal charge screening, distinct phase separated states of AL-tau complexes coexisted with liquid-liquid phase separated tau self-coacervates due to the polyampholytic nature of tau. AL-tau complex morphologies included tau-decorated giant multi- or uni-lamellar vesicles with bound tau-membrane domains (i.e. smaller tau-coated liposomes attached to the giant vesicles), and tau-mediated assemblies of smaller liposomes glued together through the cationic domains of tau. The tau self-coacervates were observed to dissolve with increasing ionic strength approaching physiological 1:1 electrolyte concentrations (100mM to 150 mM). In contrast, AL- tau complexes remained stable in the higher salt concentration conditions implying that the electrostatic interactions of tau with anionic membranes is significantly stronger than the electrostatic interactions stabilizing tau self-coacervates (i.e. where tau self-coacervates are stable only at relatively low ionic strengths much less than the average concentrations found in axons).

The findings are consistent with the hypothesis that the overall positively charged tau may interact with anionic lipid domains of the lumen-facing monolayer of the axonal plasma membrane, through long-lived or transient interactions at physiologically relevant ionic strengths. Such interactions may also contribute to the process of filopodial/growth cone outgrowth during development and in neuronal regeneration.^62,63^ For tau bound to MTs, the relevant cationic domains involved in interactions with negative lipid domains include the PRR and large sections of the C-terminal domain, which flank the MTBR region (i.e. to allow for different sections of tau to simultaneously bind MTs and negative membrane domains). For free tau, the heavily cationic MTBR is also a cite for tau-membrane interactions.

The anionic liposome-tau complexes described in this study may be referred to as tau- lipoplexes in analogy with lipoplexes consisting of cationic liposome-DNA complexes used in nucleic acid delivery applications.^43–46^ While the current study focused on elucidating phase separated structures on the optical micron scale, we expect that future small-angle X-ray scattering studies, both in bulk phases and dilute solutions similar to conditions in this study, will reveal a rich variety of distinct structures (including lamellar, hexagonal, and cubic phases) on the nanometer and subnanometer scales that have been observed for CL-nucleic acid lipoplexes.

## ACKNOWLEDGEMENTS

The authors dedicate the paper to the memory of the biophysicist Erich Sackmann. The work was supported by the Department of Energy, Office of Basic Energy Sciences, Division of Materials Sciences and Engineering, under award DE-FG02-06ER46314 (CRS and YL, charged bio- assemblies inspired by nature) and, in part, by the US National Science Foundation, Division of Materials Research, under award DMR-1807327 (CRS, phase behavior in biomaterials). SCF acknowledges support from the US National Institutes of Health grant NS-35010 and the Academic Senate of the University of California at Santa Barbara. AJJ was supported by a Research Internships in Science and Engineering (RISE) Fellowship sponsored by the Materials Research Science and Engineering Center (MRSEC) at UC Santa Barbara under award NSF DMR–2308708. CRS acknowledges sabbatical leave for Spring quarter 2024, which allowed for scholarly activities related to the research reported here.

## TOC Graphic

For Table of Contents Only

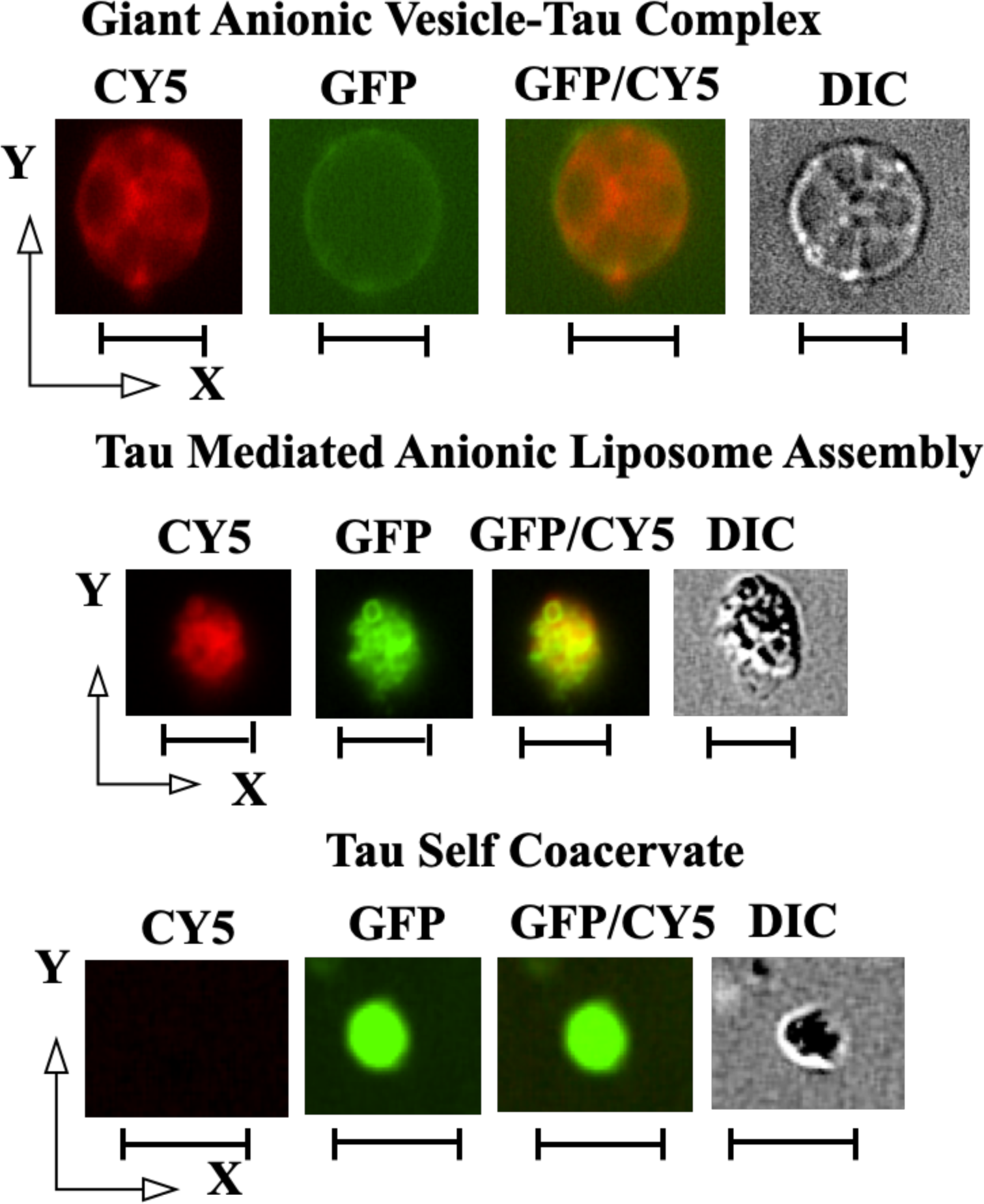

